# MPA_Pathway_Tool: User-friendly, automatic assignment of microbial community data on metabolic pathways

**DOI:** 10.1101/2021.07.07.450993

**Authors:** Daniel Walke, Kay Schallert, Prasanna Ramesh, Dirk Benndorf, Emanuel Lange, Udo Reichl, Robert Heyer

## Abstract

**Motivation:** Taxonomic and functional characterization of microbial communities from diverse environments such as the human gut or biogas plants by multi-omics methods plays an ever more important role. Researchers assign all identified genes, transcripts, or proteins to biological pathways to better understand the function of single species and microbial communities. However, due to the versatility of microbial metabolism and a still increasing number of new biological pathways, linkage to standard pathway maps such as the KEGG (Kyoto Encyclopedia of Genes and Genomes) central carbon metabolism is often problematic.

**Results:** We successfully implemented and validated a new user-friendly, stand-alone web application, the MPA_Pathway_Tool. It consists of two parts, called ‘Pathway-Creator’ and ‘Pathway-Calculator’. The ‘Pathway-Creator’ enables an easy setup of user-defined pathways with specific taxonomic constraints. The ‘Pathway-Calculator’ automatically maps microbial community data from multiple measurements on selected pathways and visualizes the results.

**Availability and Implementation:** The MPA_Pathway_Tool is implemented in Java and ReactJS. It is freely available on http://mpa-pathwaymapper.ovgu.de/. Further documentation and the complete source code are available on GitHub (https://github.com/danielwalke/MPA_Pathway_Tool).

**Supplementary Information:** Additional files and images are available at *MDPI* online.

**Highlights:** user-friendly generation of pathways, re-using of existent metabolic pathways, automated mapping of data

## Motivation

In the last years, studying the taxonomic and functional composition of bacterial, viral, and archaeal species in the human gut [1–5], in biotechnology [6–8], or the environment [9,10] became increasingly important [8]. There are several different approaches for analyzing microbial communities, focussing on the entirety of the genes (metagenomics), transcripts (metatranscriptomics), or proteins (metaproteomics). Metagenomics only reveals the presence of genes, whereas metatranscriptomics and metaproteomics indicate actual protein expression [8]. Based on the protein expression levels, a microbial communities’ phenotype can be linked with specific environmental conditions, process parameters, or diseases [11]. Due to the complexity and amount of the multi-omics data, comprehensive bioinformatic workflows were developed for the data evaluation [12–15]. For example, the MetaProteomeAnalyzer (MPA) enables analyzing and interpreting metaproteomics data sets. It offers a free, open-source, end-user-oriented complete pipeline from peak lists to taxonomic and functional result evaluation. Among others, the MPA links identified proteins to functional categories (e.g., biological keywords) and the KEGG pathways [16]. In addition to the KEGG pathway system [17], several other pathway collection and mapping tools such as Reactome [18], Pathview Web [19], Escher [20], Pathway Tools [21] exist supporting the data analysis of omics-datasets. However, due to the microbial metabolism’s versatility and constantly newly discovered biological pathways [22], linkage to standard pathway maps is insufficient for many microbial community studies. Therefore new tools are required, tailored for microbial community studies.

In general, a good pathway tool needs to meet at least the following six criteria. (i.) It should provide easy and intuitive creation of pathways to enable the fast generation of multiple pathways. (ii.). Since new reactions are discovered and pathways might be updated, the tool should support modifying the pathway maps, i.e., appending new and deleting existent reactions. (iii.) Already created pathways from different databases should be reusable. Consequently, the pathway tool should provide an import function for standard exchange formats, like comma-separated values (CSV), JavaScript Object Notations (JSON), and Systems Biology Markup Language (SBML) formats. (iv.) The pathway tool should also map experimental data on created pathways and highlight differences between the considered samples. (v.) Since metabolic reactions are taxonomy-specific, the pathway tool needs a filter to distinguish between reactions carried out by a specific taxonomy. One example of this specificity is hydrogenotrophic methanogenesis and the Wood-Ljungdahl pathway. Both pathways share similar enzymes (i.e., similar EC-numbers (Enzyme Commission numbers)). However, hydrogenotrophic methanogenesis is carried out only by archaea [23], while the Wood-Ljungdahl pathway is carried out mainly by bacteria [8]. (vi.) The tool should be independent of operating systems so that nearly everyone can use the tool, favoring an implementation as a web application.

The paper aims to present the new web application called MPA_Pathway_Tool. It enables easy creation of user-defined pathways with specific taxonomic constraints, automatically mapping microbial community data from multiple measurements on selected pathways, and visualizing the results (‘Pathway-Creator’). Additionally, the ‘Pathway-Calculator” enables mapping an entire omics data set on multiple pathways to support the automated data analyses. The functionality of the MPA_Pathway_Tool is demonstrated and validated by reproducing the manual assignment of proteins to metabolic pathways from a previous metaproteomics study about biogas plants, focussing on three pathways (hydrogenotrophic methanogenesis, acetoclastic methanogenesis, and Wood-Ljungdahl pathway).

## Results

### The ‘Pathway-Creator’ enables users to define their own metabolic pathways

The first part of the MPA_Pathway_Tool is the ‘Pathway-Creator’ (figure 1). It enables the creation of user-defined pathways by adding reactions iteratively and linking omics data to this specific pathway. The left side of the ‘Pathway-Creator’ contains a list of buttons for uploading experimental data and pathways (as CSV, JSON, and SBML), adding new reactions from KEGG, adding user-defined reactions, importing multiple reactions, and downloading created pathways (as CSV, SBML, JSON, and Scalable Vector Graphics (SVG)) and mapped data (as CSV). The right side contains a graph for visualizing the created pathway. Circular-shaped nodes are metabolites (in KEGG referred to as compounds), and diamond-shaped nodes are reactions. Nodes are connected by edges, which display the direction of a metabolic reaction. After uploading experimental data, a further user interface emerges, showing the mapping of the data set to the pathway. After sample selection, by clicking on the respective button at the bottom of the tool, reaction nodes are colored dependent on their abundance in the sample and the color settings. Information about abundances in all samples for a specific reaction is available by clicking on the respective reaction-node.

**Figure 1:**
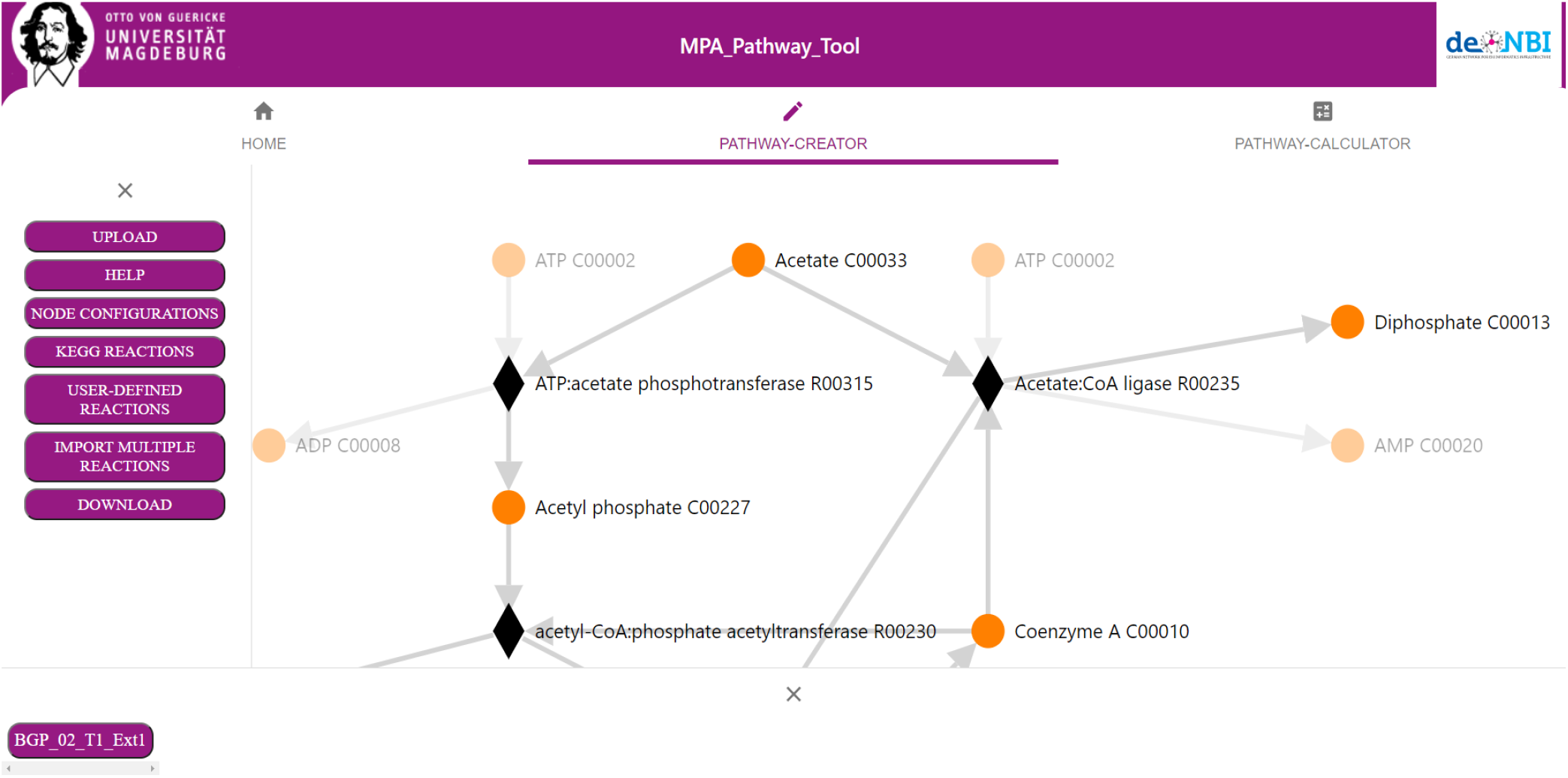
Screenshots of the ‘Pathway-Creator’ of the MPA_Pathway_Tool. The ‘Pathway-Creator’ enables the creation of user-defined pathways by adding reactions iteratively. A user-interface (left side) allows users to upload experimental data and pathways, add new reactions, modify the pathway’s visualization, and download the created pathways and the mapped data. A network graph (right side) visualizes the created pathway. Metabolites are visualized as circular nodes, while reactions are visualized as diamond-shaped nodes. The connecting edges show the direction of each reaction. On the bottom are buttons for selecting samples of the experimental data and a heatmap for adjusting the colors. Users can choose a sample, and as a consequence, the reaction nodes are colored according to their abundance.

### The ‘Pathway-Calculator’ enables automated mapping of experimental data on multiple metabolic pathways

The ‘Pathway-Calculator’ (figure 2) consists of two upload zones, one for experimental data and another for multiple pathway files (as CSV, JSON, or SBML). The ‘Pathway-Calculator’ performs mapping of experimental data on multiple uploaded pathways. The mapping algorithm is explicitly explained in methods. After uploading all previously created pathways (CSV) and the experimental data in the ‘Pathway-Calculator’, the calculation starts. Subsequently, the result table can be exported as CSV. Furthermore, a list with all unmatched features (e.g., proteins) can be downloaded.

**Figure 2:**
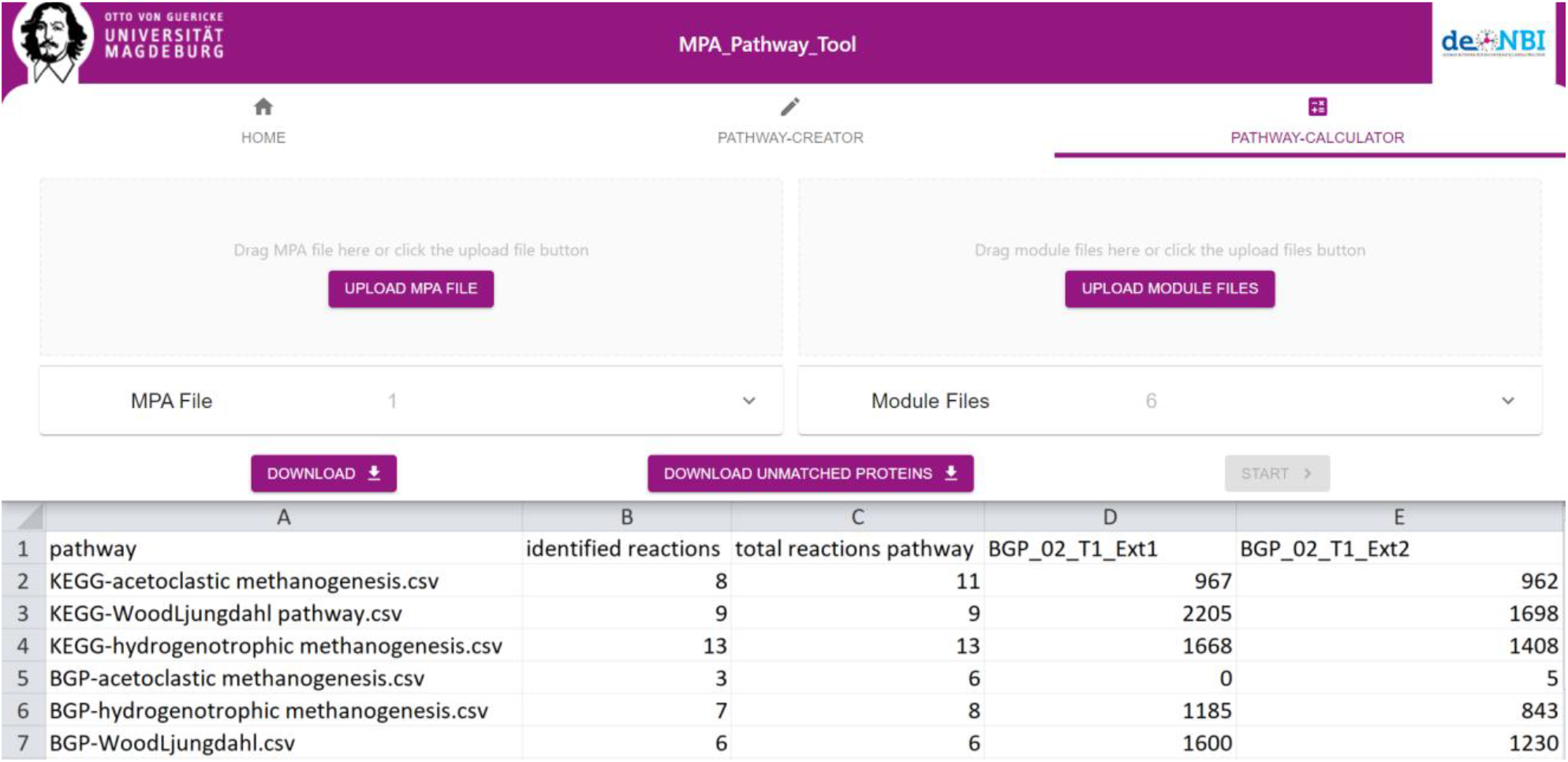
Screenshot of the ‘Pathway-Calculator’ of the MPA_Pathway_Tool. Users can upload experimental data as CSV on the left and multiple created pathways (CSV, SBML, and JSON) on the right side. After starting by clicking ‘start’, the ‘Pathway-Calculator’ matches the features (e.g., proteins) on all pathways and returns a file with the results (for further details, refer to the mapping algorithm in the material and methods part). Additionally, users can download a file of all unmatched features (e.g., proteins).

Finally, we tested the performance of the ‘Pathway-Calculator’ by uploading experimental data with different file sizes (10,000 proteins, 100.000 proteins, and 1,000,000 proteins with 44 samples per test) and different a number of pathways on one of our local desktop computers (AMD Ryzen 5 3600, 16 GB DDR4 RAM 3000MHz, Chrome Browser version 89.0.4389.90). We utilized 1, 10, and 100 copies of the user-defined Wood-Ljungdahl pathway used in Heyer et al. 2019b (Biogas plant (BGP)-Wood-Ljungdahl pathway) for the performance test. Finally, we measured the elapsed time between pressing the ‘start’-button and the availability of the download link (table 1). Each test with 10,000 and 100,000 proteins performed under 1 minute indicating a good performance for most files. Tests with 1,000,000 proteins took much longer (up to 12 minutes) caused by higher upload times and high requirements on memory and on CPU performance (table 1).

**Table 1:**
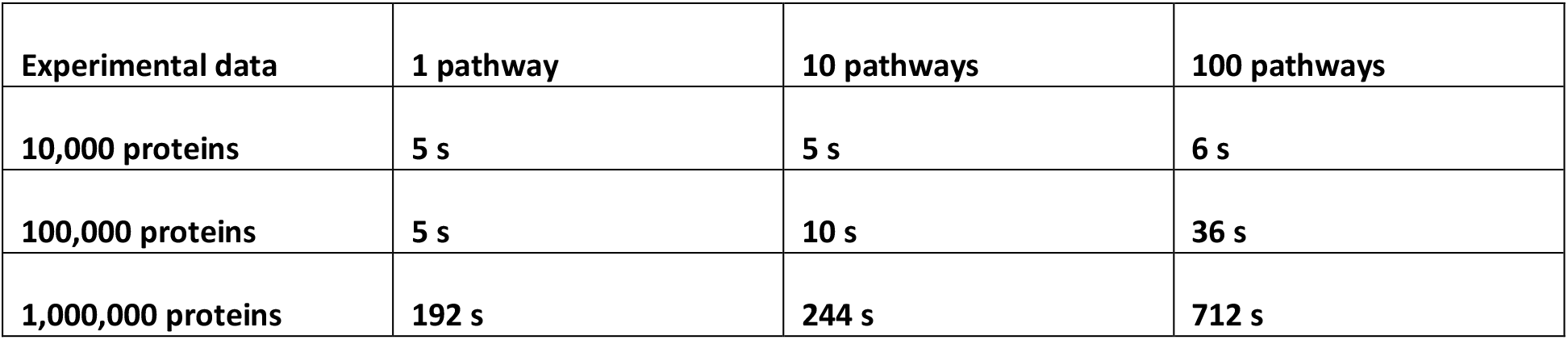
Performance test for the ‘Pathway-Calculator’. It lists the elapsed time in seconds between pressing ‘start’ and receiving the download link (AMD Ryzen 5 3600, 16 GB DDR4 RAM 3000MHz, Chrome version 89.0.4389.90) Each test was performed with 44 samples and different numbers of proteins.

| Experimental data | 1 pathway | 10 pathways | 100 pathways |
| --- | --- | --- | --- |
| 10,000 proteins | 5 s | 5 s | 6 s |
| 100,000 proteins | 5 s | 10 s | 36 s |
| 1,000,000 proteins | 192 s | 244 s | 712 s |

### Tool validation with experimental data

Validation of the MPA_Pathway_Tool was carried out by comparing our pathway assignment against a manual pathway assignment of a previous metaproteomics study about biogas plants [8]. Additionally, we assigned the proteins to the standard KEGG modules to illuminate the demand for user-specific pathways.

For the evaluation, we created six pathway maps (BGP-hydrogenotrophic-methanogenesis, BGP-acetoclastic-methanogenesis, BGP-Wood-Ljungdahl-pathway, KEGG-hydrogenotrophic-methanogenesis, KEGG-acetoclastic-methanogenesis, and KEGG-Wood-Ljungdahl-pathway). The three user-defined pathways (BGP-, Supplementary information figure 3-5) were created by importing EC-numbers under ‘import multiple reactions’ and selecting the appropriate reactions (for further details, see the tutorial in supplementary information file 17) based on the pathway assignment in [8]. The creation of each of the other three KEGG pathways (Supplementary information figure 6-8) was performed by importing the respective KEGG-MODULE using ‘import KEGG MODULE’ under ‘import multiple reactions’. For each pathway, a taxonomic classification was added (table 2) by using ‘Add Taxonomy’ under ‘node configurations’ and adding the respective taxonomy. According to the publication of the dataset, we decided to exclude Archaea from the Wood-Ljungdahl pathway and include only Archaea for the hydrogenotrophic and acetoclastic methanogenesis [8]. Subsequently, the experimental metaproteomics data (Supplementary information file 9) were mapped on each pathway using single pathway (Supplementary information file 10-15) and multiple pathway mapping (figure 4)(Supplementary information file 1). Unmatched proteins were downloaded as CSV (Supplementary information file 2).

**Table 2:** Taxonomic classification of each pathway. Taxonomic assignments for the BGP-pathways were carried out according to the manual assignments in the used data set for the validation.

| Pathway | Added taxonomic requirement |
| --- | --- |
| BGP-Hydrogenotrophic methanogenesis (user-defined) | only Archaea |
| KEGG-Hydrogenotrophic methanogenesis (KEGG-MODULE) | only Archaea |
| BGP-Acetoclastic Methanogenesis (user-defined) | only Archaea |
| KEGG-Acetoclastic Methanogenesis (KEGG-MODULE) | only Archaea |
| BGP-Wood- Ljungdahl pathway (user-defined) | all except Archaea |
| KEGG-Wood- Ljungdahl pathway (KEGG-MODULE) | all except Archaea |

**Figure 3:**
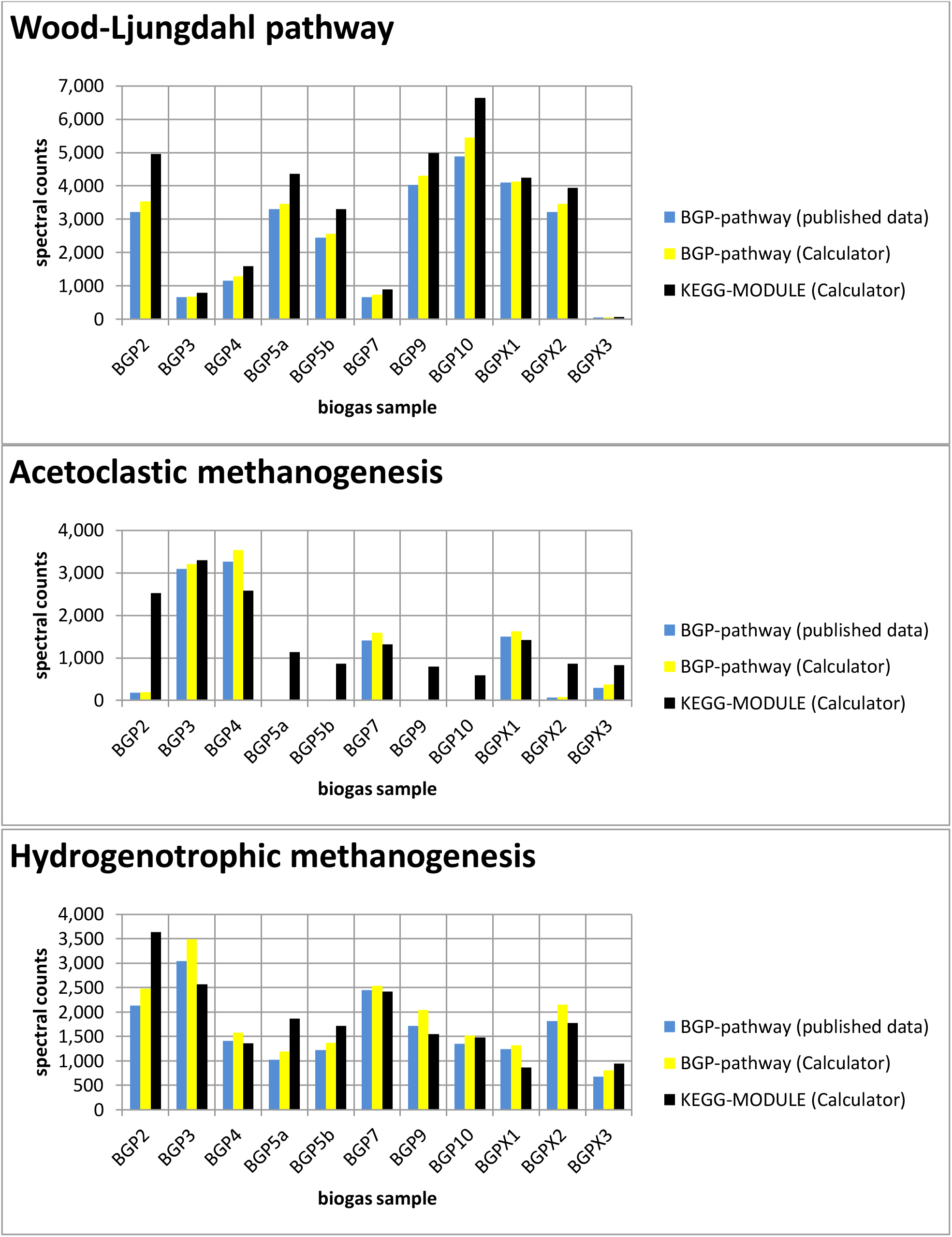
Comparison of experimental data mapped on each pathway (Wood-Ljungdahl pathway, acetoclastic methanogenesis, and hydrogenotrophic methanogenesis) with published data [8]. Summed spectral counts plotted against all biogas samples are blue and yellow for user-defined pathways and black for KEGG-MODULEs. The results were obtained from the ‘Pathway-Calculator’ (yellow and black) and published data (blue). Especially acetoclastic methanogenesis shows higher spectral counts in most samples, indicating the occurrence of unspecific reactions.

**Figure 4:**
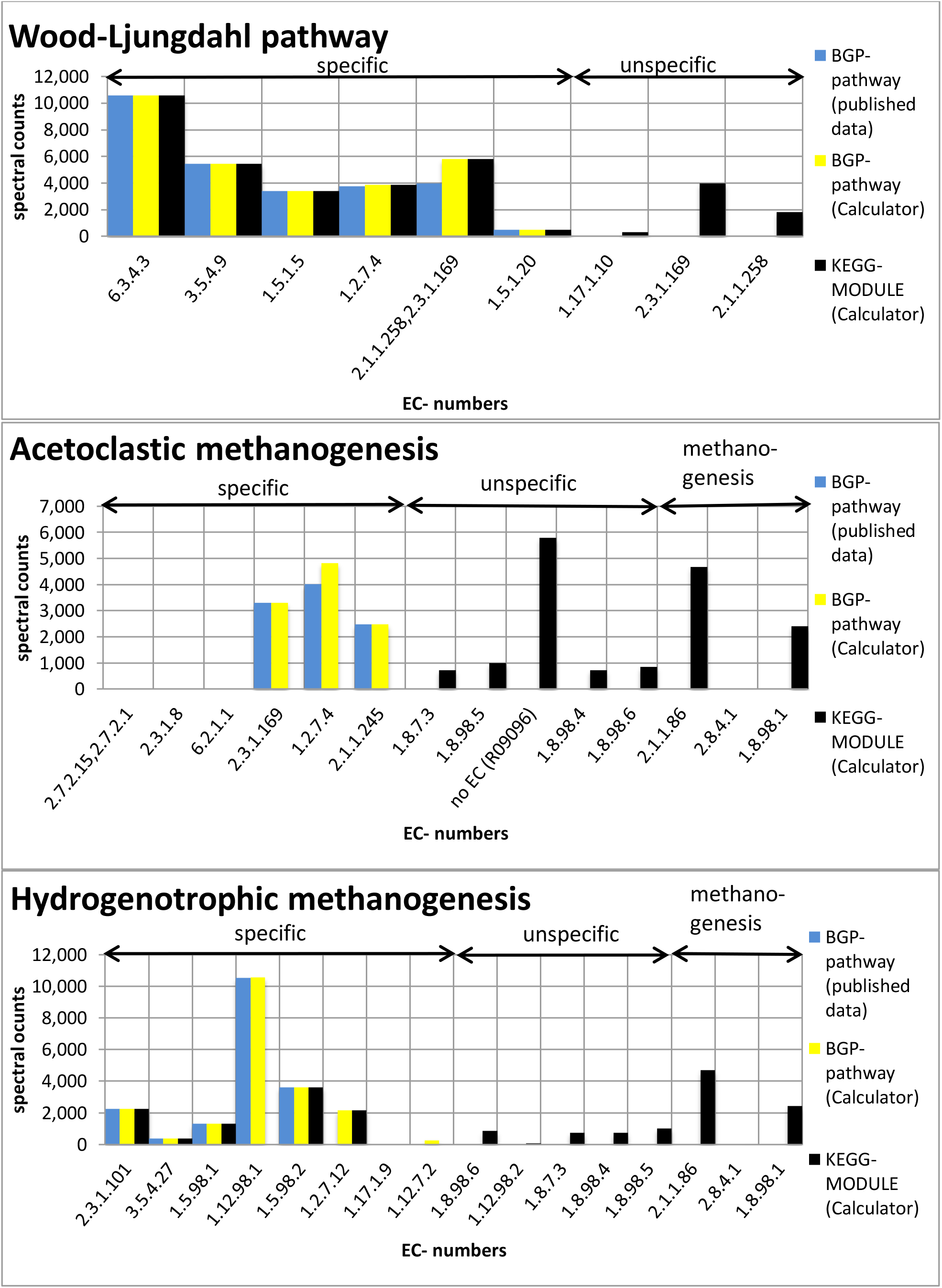
Summed spectral counts of each reaction in the Wood-Ljungdahl pathway, acetoclastic methanogenesis, and hydrogenotrophic methanogenesis. Results from published data [8] were visualized in blue, results from the ‘Pathway-Calculator’ for the user-defined in yellow, and results from the ‘Pathway-Calculator’ for KEGG-MODULEs in black. All KEGG-MODULEs possess additional unspecific reactions. KEGG-acetoclastic-methanogenesis and KEGG-hydrogenotrophic methanogenesis possess three additional reactions (2.1.1.86, 2.8.4.1., and 1.8.98.1), which catalyze the last steps of the methanogenesis. (6.3.4.3: Formate:tetrahydrofolate ligase (ADP-forming); 3.5.4.9: 5,10-Methenyltetrahydrofolate 5-hydrolase; 1.5.1.5: 5,10-methylenetetrahydrofolate:NADP+ oxidoreductase; 1.2.7.4: carbon-monoxide,water:ferredoxin oxidoreductase; 2.1.1.258,2.3.1.169: Tetrahydrofolate + Acetyl-CoA <=> 5-Methyltetrahydrofolate + CoA + CO; 1.5.1.20: 5-methyltetrahydrofolate:NADP+ oxidoreductase; 1.17.1.10: formate:NADP+ oxidoreductase; 2.3.1.169: acetyl-CoA:corrinoid protein O-acetyltransferase; 2.1.1.258: 5-methyltetrahydrofolate:corrinoid/iron-sulfur protein methyltransferase; 2.3.1.101: Formylmethanofuran:5,6,7,8-tetrahydromethanopterin 5-formyltransferase; 3.5.4.27: 5,10-Methenyltetrahydromethanopterin 10-hydrolase; 1.5.98.1: 5,10-methylenetetrahydromethanopterin:coenzyme-F420 oxidoreductase; 1.12.98.1: Hydrogen:Coenzyme F420 oxidoreductase; 1.5.98.2: 5,10-Methylenetetrahydromethanopterin:coenzyme-F420 oxidoreductase; 1.2.7.12: formylmethanofuran:ferredoxin oxidoreductase; 1.17.1.9: formate:NAD+oxidoreductase; 1.12.7.2: hydrogen:ferredoxin oxidoreductase; 1.8.98.6: coenzyme B,coenzyme M,ferredoxin:formate oxidoreductase; 1.12.98.2: hydrogen:N5,N10-methenyltetraydromethanopterin oxidoreductase; 2.1.1.86: 5-methyl-5,6,7,8-tetrahydromethanopterin:2-mercaptoethanesulfonate 2-methyltransferase; 1.8.7.3: CoB,CoM:ferredoxin oxidoreductase; 1.8.98.4: CoB,CoM,ferredoxin:coenzyme F420 oxidoreductase; 2.8.4.1: 2-(methylthio)ethanesulfonate:N-(7-thioheptanoyl)-3-O-phosphothreonine S-(2-sulfoethyl)thiotransferase; 1.8.98.5: CoB,CoM,ferredoxin:H2 oxidoreductase; 1.8.98.1: Coenzyme B:coenzyme M:methanophenazine oxidoreductase; 2.7.2.15,2.7.2.1: ATP:acetate phosphotransferase; 2.3.1.8: acetyl-CoA:phosphate acetyltransferase; 6.2.1.1: Acetate:CoA ligase; 2.1.1.245: 5-methyltetrahydrosarcinapterin:corrinoid/iron-sulfur protein methyltransferase, R09096: acetyl-CoA decarbonylase/synthase)

**Figure 5:**
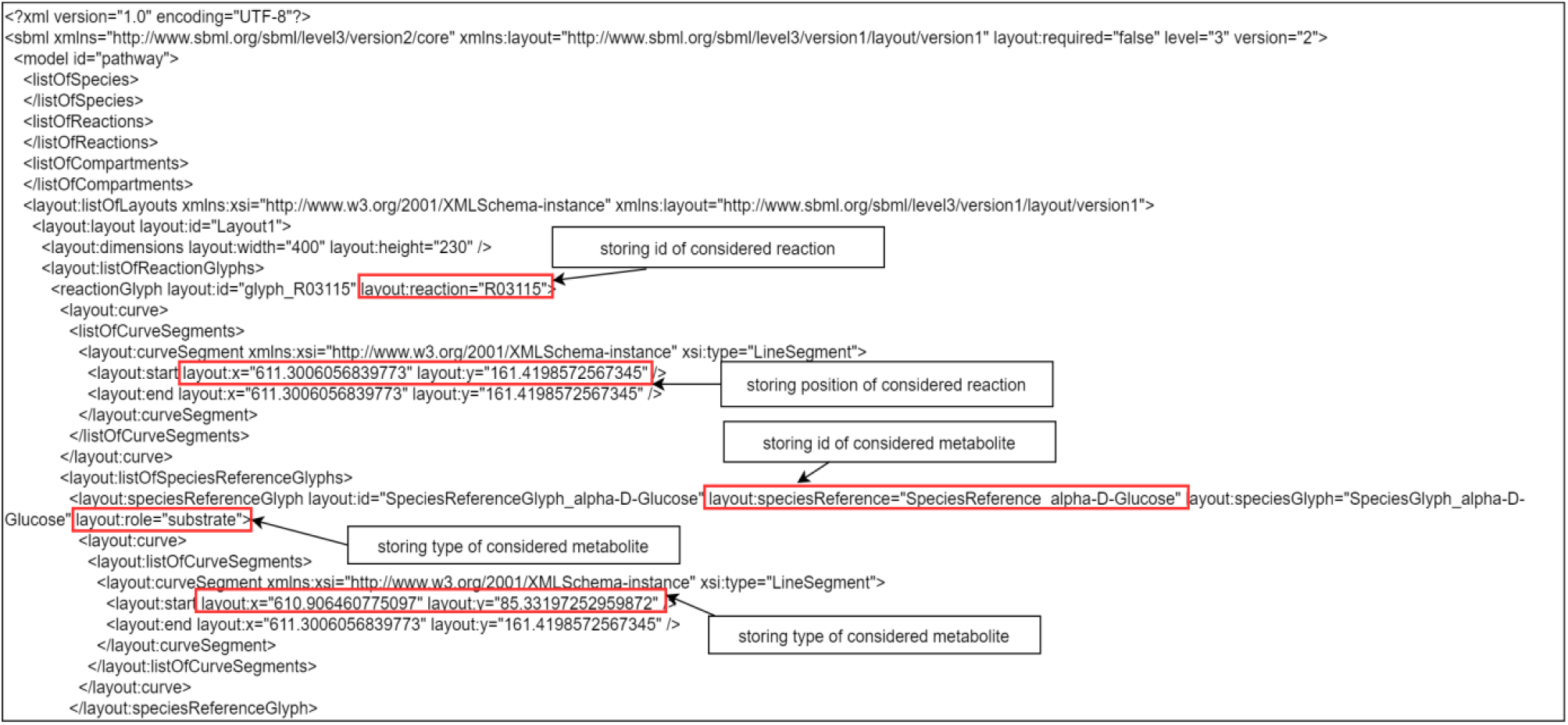
An example of updates in an SBML-file. In the tag ‘layout:listOfLayouts’ all positions for the nodes of the created pathway are stored.

**Figure 6:**
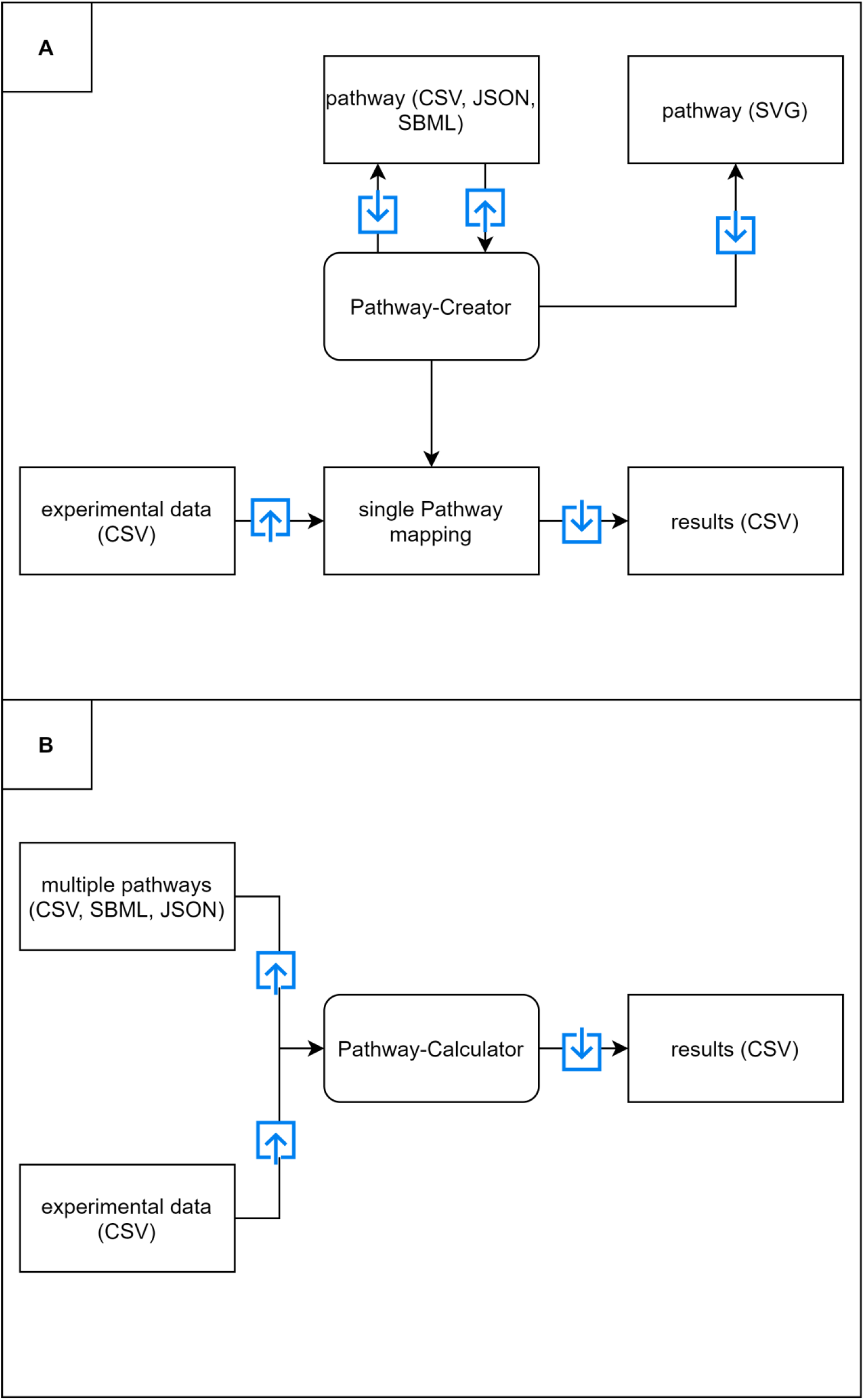
An overview of the structure of the MPA_Pathway_Tool with the ‘Pathway-Creator’ (A) and the ‘Pathway-Calculator’ (B). A) The ‘Pathway-Creator’ provides the creation of user-defined pathways. These pathways can be exported as SBML, SVG, CSV, or JSON. For later modifications, the user can also upload these pathways (CSV, JSON, SBML). Furthermore, the ‘Pathway-Creator’ enables the mapping of experimental data (CSV) on the created pathway. Results can be exported as CSV. B) The ‘Pathway-Calculator’ provides the mapping of experimental data (CSV) on multiple pathways (CSV, JSON, and SBML). The resulting mapped data can also be exported as CSV. Blue arrows represent an upload or download of specific files.

The summed spectral counts mapped on each user-defined pathway were slightly greater than in the published experimental data for each sample. The observed differences had three causes. First, some reactions have multiple EC-numbers. In the published experimental data, only one EC-number was considered for each reaction. The second reason for the differences is the missing matching of proteins by K-numbers, which also catalyze the same reaction. They were not considered in the published data. The last reason is the usage of deprecated EC-numbers in the published data. EC-numbers are transferred to new EC-numbers occasionally. All three reasons lead to a minor loss of information in the published dataset due to less matched proteins. The stated issues were corrected in the MPA_Pathway_Tool, resulting in a higher accuracy during the pathway assignment.

Furthermore, we compared the results of each BGP-pathway with the corresponding KEGG-pathway. We observed higher spectral counts for the created KEGG MODULEs for most analyzed pathways compared to the respective BGP-pathways (figure 3). Especially acetoclastic methanogenesis exhibits higher spectral counts in most samples for the KEGG MODULE. One reason for higher spectral counts in KEGG MODULEs is the integration of more reactions in the KEGG MODULEs than in the BGP-pathways (figure 4). Whereas KEGG modules contain all reactions for a certain function, BGP pathways focus on the pathway specific for the function. For example, KEGG-acetoclastic-methanogenesis, and KEGG-hydrogenotrophic methanogenesis possess three additional reactions (2.1.1.86, 2.8.4.1., and 1.8.98.1), which catalyze the last steps of the methanogenesis. These were purposely integrated into another separate BGP-pathway, because of the occurrence in hydrogenotrophic and acetoclastic methanogenesis. This separation prevented repeated summation of spectral counts of equal reactions. The analyzed KEGG MODULEs additionally possess similar reactions with minor differences (e.g., NADH+H^+^ or NADPH+H^+^ as reductant).

### Conclusion: The MPA_Pathway_Tool provides an easy and fast option to set up multiple pathways

We successfully implemented the pathway tool to meet all of our six defined criteria. (i.) The MPA_Pathway_Tool provides an easy and fast setup of multiple pathways. Multiple reactions can be imported using various options, e.g., import by EC numbers, import of a KEGG MODULE, or entire SBML files. (ii.) A further modification of the generated pathways is possible by deleting reactions and adding new reactions (from the KEGG database and user-defined reactions). (iii.) As interchange formats, JSON, CSV, and SBML were implemented. (iv.) Experimental data from metaproteomics, metatranscriptomics, and metagenomics studies can be automatically mapped on single pathways (‘Pathway-Creator’) and multiple pathways (‘Pathway-Calculator’). The results of pathway mapping can also be exported as CSV. (v.) The mapping algorithm includes a taxonomic filter, which was successfully applied by comparing our results with published experimental data for the stated pathways. (vi.) The MPA_Pathway_Tool was implemented as a stand-alone web application to guarantee the independence from users’ operating systems.

### Future work: integration to other features and addition of new functions will increase the flexibility of the MPA_Pathway_Tool

The MPA_Pathway_Tool is developed as a stand-alone application for the characterization and analysis of microbial community data. To broaden its scope of application, it will be integrated into the next version of the MPA software. However, in the future other tools such as UniPept [24] or Prophane [25] might be linked, too.

Since the MPA_Pathway_Tool already provides stoichiometric data, further options like identification-driven flux balance analysis and flux vector analysis might be added to the workflow using tools such as the CellNetAnalyzer [26], COBRApy [27] or Escher [20]. As it allows the calculation of metabolic flows in metabolic networks, it can be used to predict the specific growth rate of organisms and the specific production rate of certain metabolites. Accordingly, identification of strategies for optimizing biotechnological processes might be feasible [28].

## Availability and Implementation

### General workflow

The MPA_Pathway_Tool represents an intuitive web application for mapping (meta)-proteome and other omics data on metabolic pathways and the creation and modification of pathways. It consists of two different parts. First, the ‘Pathway-Creator” allows the creation and modification of pathways. The ‘Pathway-Creator’ also supports single pathway mapping. The second part is the ‘Pathway-Calculator’, which provides multiple pathway mapping.

### Implementation

The MPA_Pathway_Tool is a stand-alone web application consisting of JavaScript (JS), Hypertext Markup Language 5 (HTML), and Cascading Style Sheets (CSS) on the client-side and Java on the server-side. The client-side is built with the help of ReactJS. We used Create React App for the setup of the project. Create React App is used to create single-page React applications, and it provides a setup without configurations [29]. Third-party dependencies have been pulled by using Node Package Manager (npm) (version 6.14.9). The most important dependencies are ReactJS (version 16.8.6) for using JSX syntax in our project, Redux (version 4.0.5) [30] and mobx (version 7.0.5) [31]-used for storing and handling states, React-d3-graph (version 2.5.0)-used for visualization of pathways [32], Axios (version 0.21.0) [33]-used for making HTTP requests from the browser, Material-UI (version 4.11.2) [34] used for the implementation of some user-interface components, Lodash (version 4.17.21) [35] for deep cloning objects, and File-saver (version 2.0.5) [36] used for saving files.

A complete list of used dependencies on the client-side is listed in the package.json file (https://github.com/danielwalke/MPA_Pathway_Tool/tree/main/keggcalculator-frontend).

We have implemented a REST-API (figure 7) and an algorithm for mapping data on multiple pathways in the programming language Java (Java SE-1.8). Dependencies have been imported using Maven. We used Gson (version 2.8.5) [37] for converting Java objects to JSON and vice versa, Spark (version 2.6.0) for setting up the REST-API, and JSBML (version 1.5) [38] for reading SBML-files.

**Figure 7:**
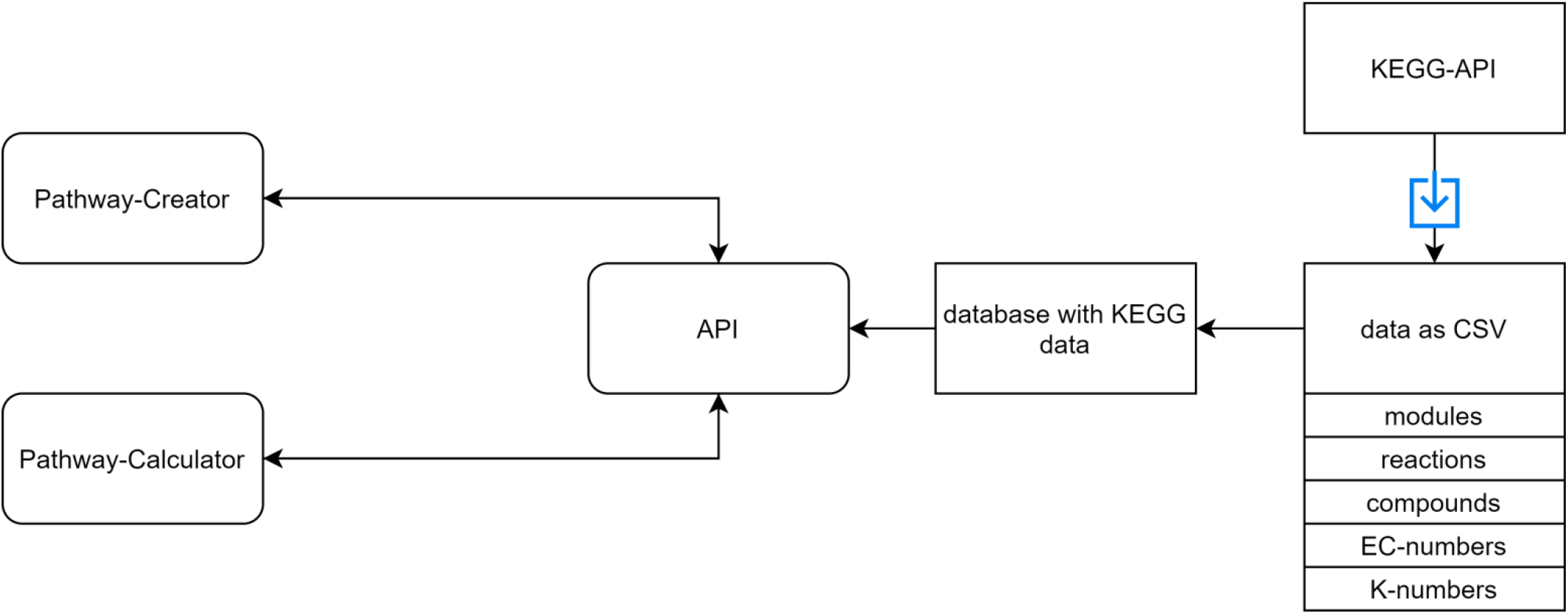
An overview of the REST-API. A REST-API is used for sending the required data to the ‘Pathway-Creator’ and the ‘Pathway-Calculator’. The KEGG database has been downloaded as CSV files and parsed into our database. The available KEGG data includes pathway modules, compounds, reactions, EC numbers, and K-numbers.

### MPA_Pathway_Tool: ‘Pathway-Creator’

In the ‘Pathway-Creator’ compounds, i.e., substrates and products (in KEGG referred to as compounds), are visualized as circular nodes and reactions as diamond-shaped nodes. Links connect compounds and reactions, and arrows show the direction of a reaction. Users have the opportunity to create their pathway from scratch by defining an initial substrate and then adding a reaction with the desired product iteratively. For each added reaction, the additional substrates and products are added automatically. The reaction library contains all reactions from the KEGG database and may extend by user-defined reactions. Multiple reactions can also be imported by using the KEGG-MODULE import, EC-number reactions import, K-number reactions import, and SBML-files. Created pathways can be exported as CSV, SBML, and JSON. In addition, to the metabolic reactions, we store in these files the opacity of nodes, their position, and the abbreviations for later usage or modifications of the pathways. This information is stored in a specific tag (‘layout:listOfLayouts’) of SBML-files (figure 5). Users can also download the created pathway as an image (SVG). Figure 6A provides an overview of the ‘Pathway-Creator’. For a detailed description of pathway creation, please refer to the tutorial in supplementary file 17.

Identified features (e.g., proteins) can be mapped on the created pathway (figure 6A). Colors visualize differences in abundances for a chosen sample for each reaction (single pathway mapping). The results can be downloaded as CSV.

### MPA_Pathway_Tool: ‘Pathway-Calculator’

Multiple created pathways (CSV, JSON, or SBML) and a file with experimental data (e.g., protein identifications) as CSV can be uploaded on the ‘Pathway-Calculator’ to perform multiple pathways mapping (figure 6B). After calculation, the result file can be downloaded as CSV, which contains in each row a pathway name, the total number of reactions in this pathway, the number of reactions identified in the data, and the summed quantified values for each sample.

### Mapping Algorithm

For a better understanding of the mapping algorithm, we explain the application with experimental data containing proteins. Of course, the mapping algorithm can be applied for other omics-data too. Each pathway contains a pathway name (filename) and multiple reactions. On the other hand, each reaction contains an ID, a name, substrates, and products with stoichiometric coefficients, EC-numbers, K-numbers, taxonomic constraints, and a list of matching proteins, which is empty at the beginning of the mapping calculation. For a better overview, only the last four properties are visualized as a scheme (figure 8). The imported file with protein identifications contains multiple proteins, which comprise an identifier, EC-numbers, K-numbers, a taxonomic tree (superkingdom, kingdom, phylum, class, order, family, genus, species), a description, and quantified values for each measured sample (Supplementary File 16, example input). For each reaction in a pathway, all proteins are analyzed. To add a protein to the list of matched proteins, two conditions must be fulfilled. The first condition is fulfilled if the EC-numbers or K-numbers of the reaction contain at least one EC-number or K-number of the protein. The second condition is fulfilled if the taxonomic tree of the protein contains the taxonomic requirement specified for the reaction. The fulfilment of both conditions adds the protein to the list of matched proteins. The quantitative values of all proteins are summed up for each sample. For single pathway mapping, the results can be visualized on the created pathway and exported as CSV. For multiple pathway mapping, all quantified values of each reaction (sums of quantified values of matched proteins) are summed up. Additionally, each pathway’s completeness is checked by returning the number of reactions identified in the data and the total number of reactions in the pathway. Finally, results can be exported as CSV. Multiple pathway mapping also provides the calculation of multiple imported pathways.

**Figure 8:**
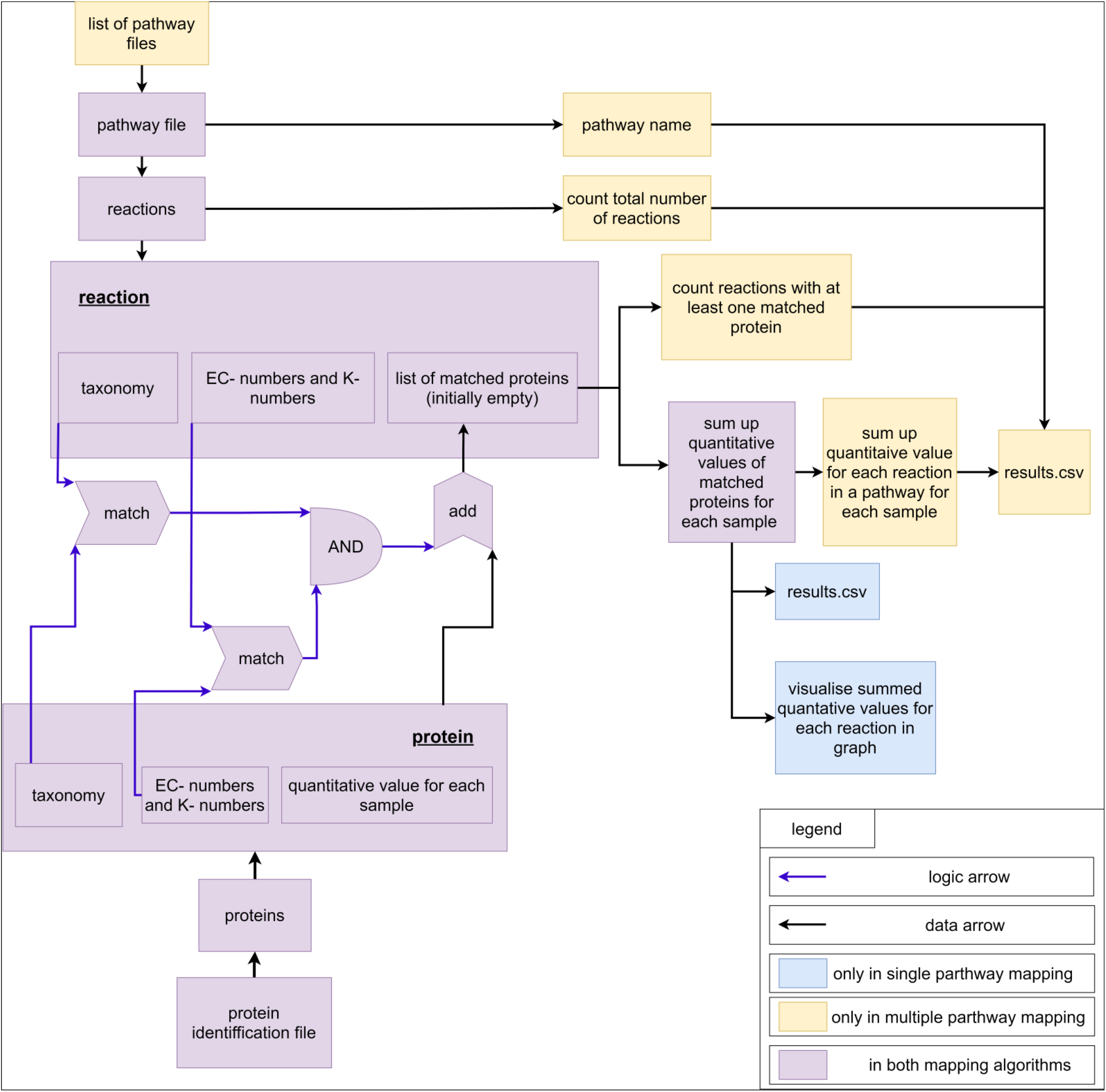
Scheme for visualization of the Mapping Algorithm.

### Experimental data

Experimental data used for validating the MPA_Pathway_Tool originate from a previous metaproteomics study of 11 different biogas plants sampled at two different time points [8]. As input, we used the final result matrix of all identified metaproteins (see Supplementary information file 5, in [8])

## Supporting information

Supplementary information file 1_result table Calculator

Supplementary information file 2_unmatched Proteins

Supplementary information file 3_BGP-WoodLjungdahl pathway

Supplementary information file 4_BGP-hydrogenotrophic methanogenesis

Supplementary information file 5_BGP-acetoclastic methanogenesis

Supplementary information file 6_KEGG-Wood-Ljungdahl pathway

Supplementary information file 7_KEGG-hydrogenotrophic methanogenesis

Supplementary information file 8_KEGG-acetoclastic methanogenesis

Supplementary information file 9_Metaprotemics data

Supplementary information file10_BGP-Wood-Ljungdahl single pathway mapping

Supplementary information file 11_BGP-hydrogenotrophic Methanogenesis single pathway mapping

Supplementary information file 12_BGP-acetoclastic Methanogenesis single pathway mapping

Supplementary information file 13_KEGG-Wood-Lungdahl single pathway mapping

Supplementary information file 14_KEGG-hydrogenotrophic Methanogenesis single pathway mapping

Supplementary information file 15_KEGG-acetoclastic methanogenesis single pathway mapping

Supplementary information file 16_format for experimental data

Supplementary information file 17_Tutorial MPA_Pathway_Tool

## Availability

The MPA_Pathway_Tool is freely available on the web at http://mpa-pathwaymapper.ovgu.de/. Further documentation and the complete source code are deposited on GitHub (https://github.com/danielwalke/MPA_Pathway_Tool).

## Funding and acknowledgment

This work was supported by the German Federal Ministry of Education and Research (de.NBI network. project MetaProtServ. grant no. 031L0103). We highly appreciate their funding.

## Supplementary information

## index of abbreviations

CSV: comma-separated values
JSON: JavaScript Object Notation
SBML: Systems Biology Markup Langugage
SVG: Scalable Vector Graphics
MPA: MetaProteomeAnalyzer
CSS: Cascading Style Sheets
HTML: Hypertext Markup Language
JS: JavaScript
EC-number: Enzyme Commission number
BGP: biogas plant
KEGG: Kyoto Encyclopedia of Genes and Genomes

