## Supplementary figures and images for "MPA_Pathway_Tool: User-friendly, automatic assignment of microbial community data on metabolic pathways"

### Supplementary information file 3_BGP-WoodLjungdahl pathway

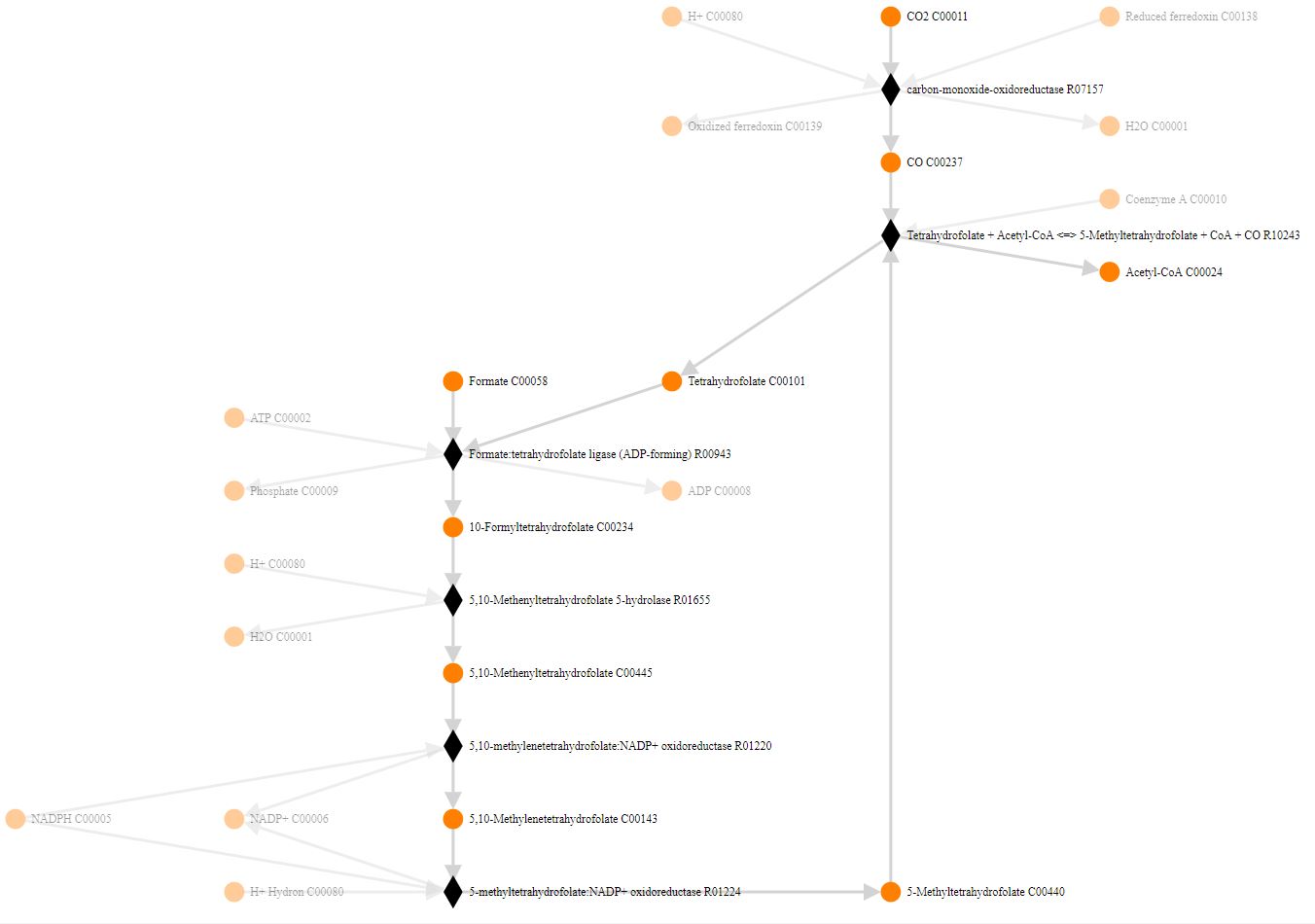

### Supplementary information file 4_BGP-hydrogenotrophic methanogenesis

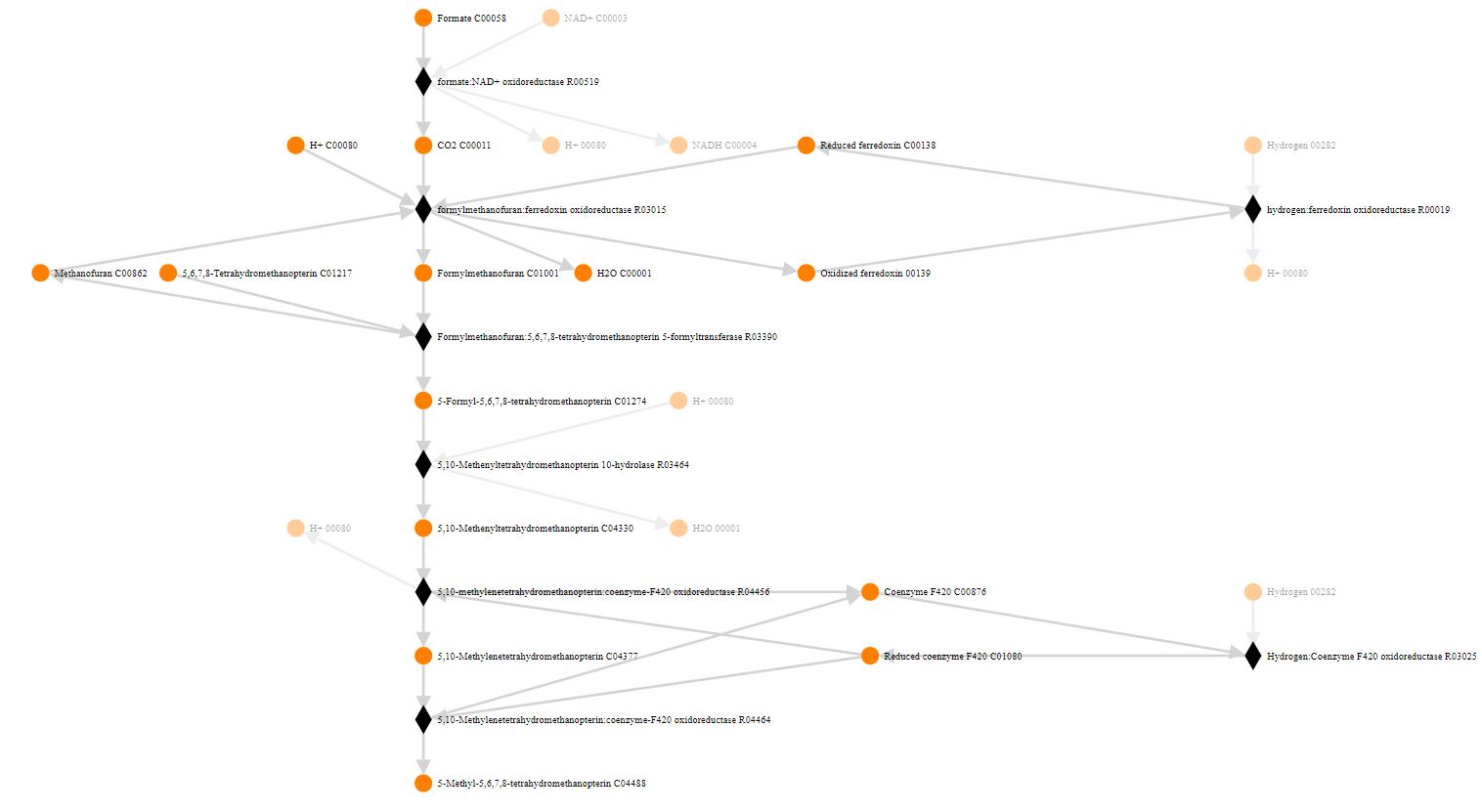

### Supplementary information file 5_BGP-acetoclastic methanogenesis

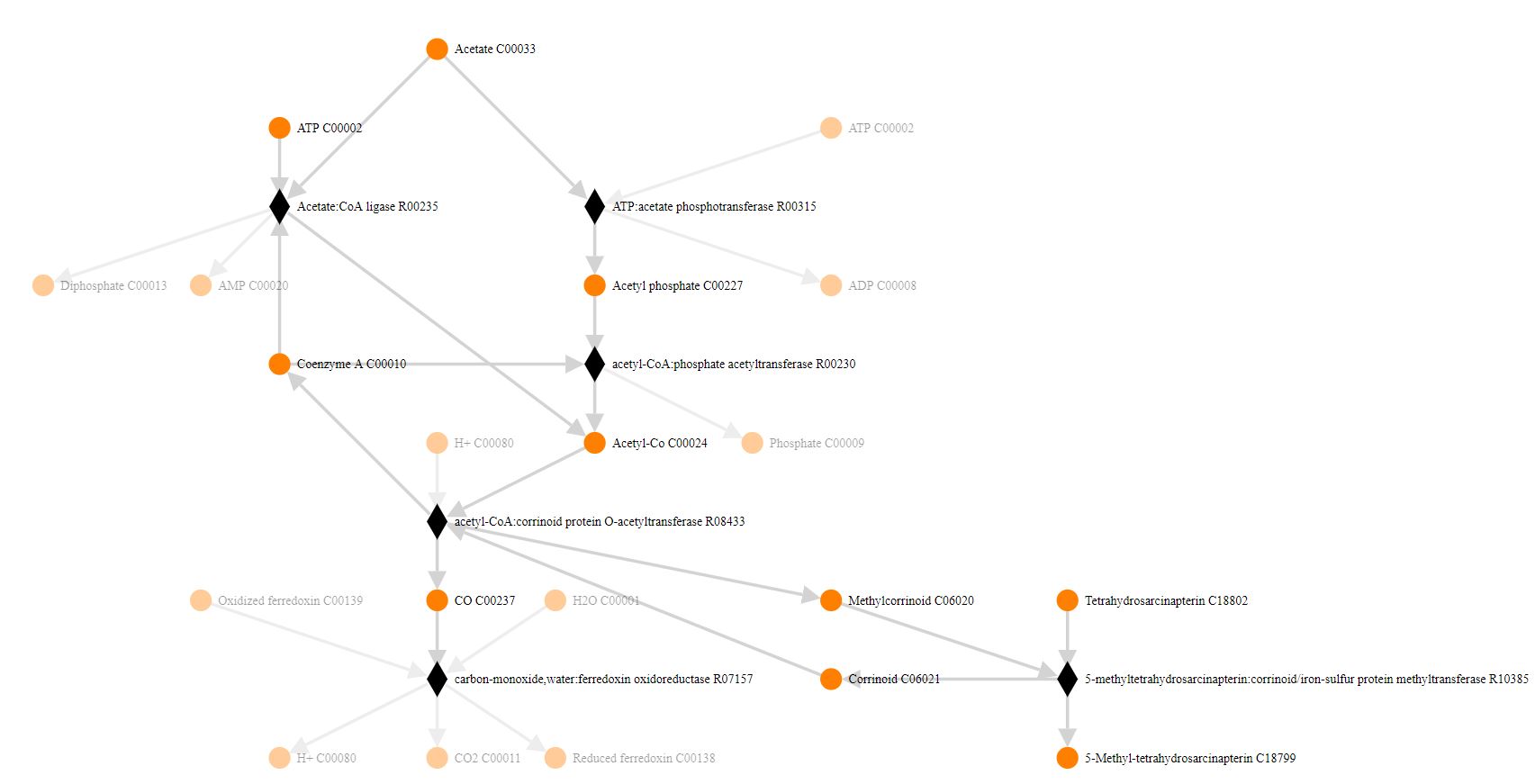

### Supplementary information file 6_KEGG-Wood-Ljungdahl pathway

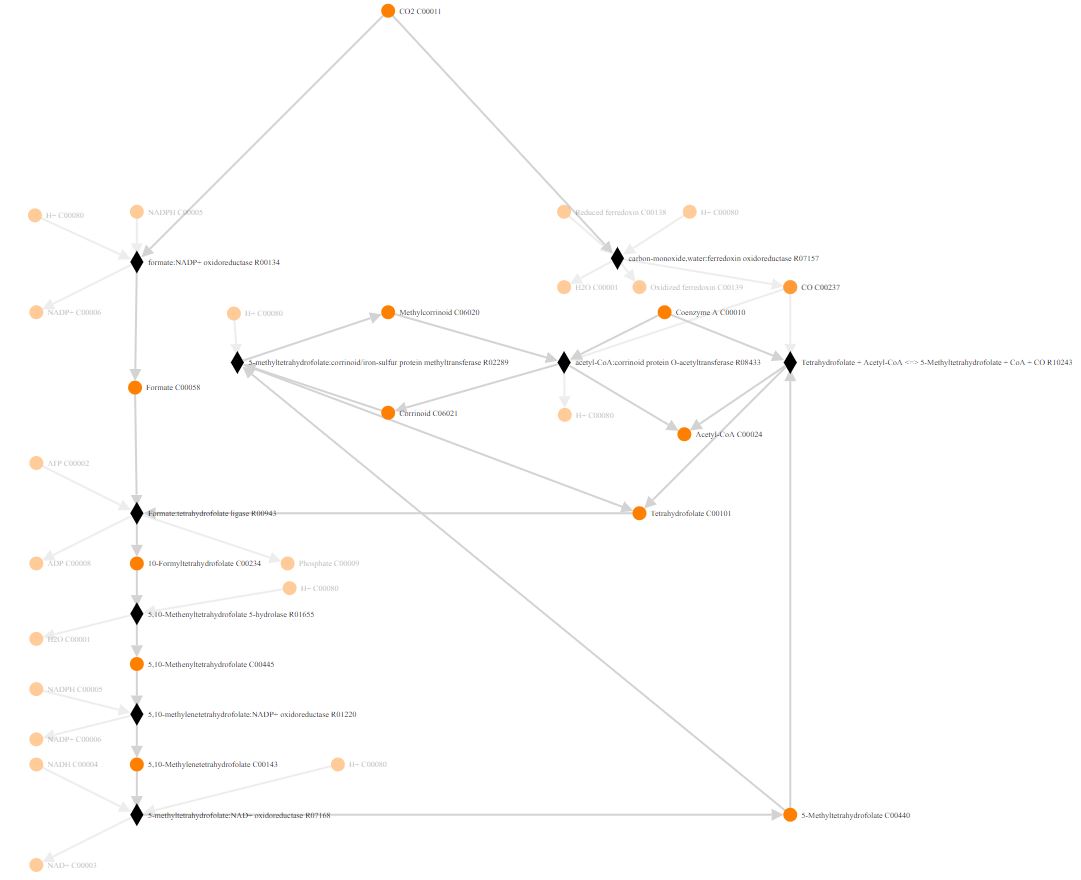

### Supplementary information file 7_KEGG-hydrogenotrophic methanogenesis

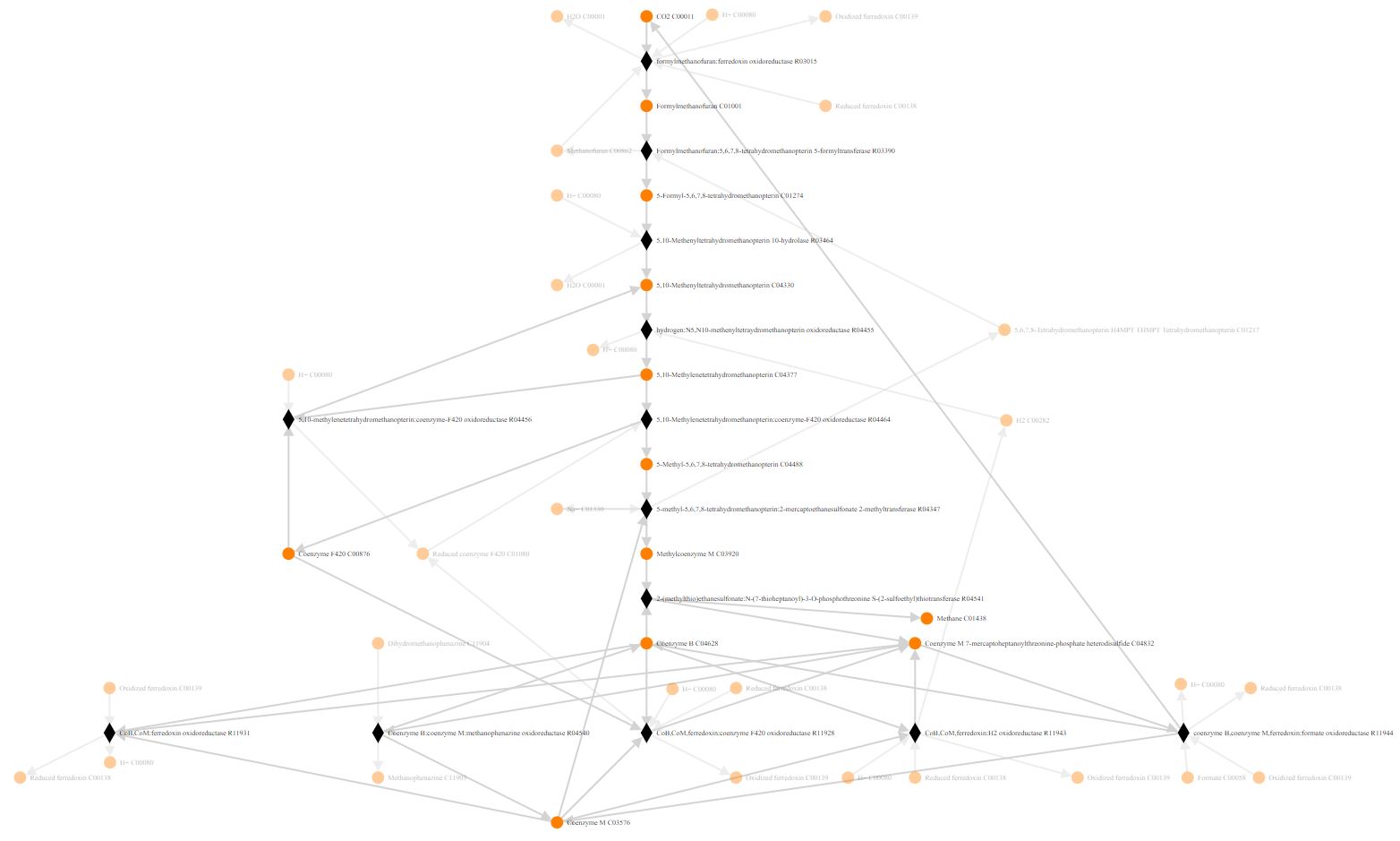

### Supplementary information file 8_KEGG-acetoclastic methanogenesis

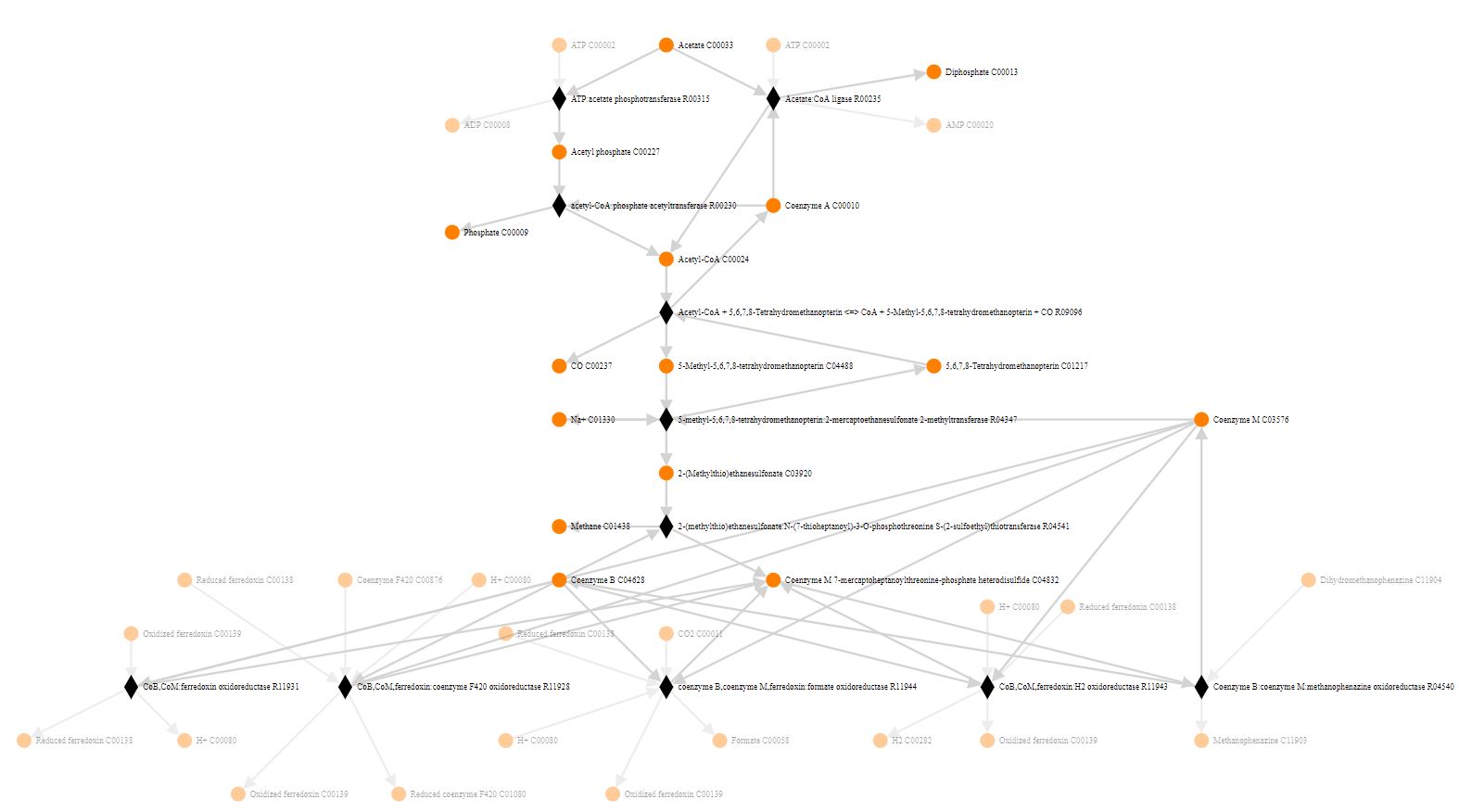
