## Supplementary information file 17_Tutorial MPA_Pathway_Tool for "MPA_Pathway_Tool: User-friendly, automatic assignment of microbial community data on metabolic pathways"

The MPA_Pathway_Tool is a new **stand-alone web application** for creating your own metabolic pathways and to map experimental data on them. The **“Pathway-Creator”** enables the **creation of metabolic pathways**, modification of existent pathways and mapping of experimental data on a single pathway. Metabolites (in KEGG referred as compounds) are visualized as circular o nodes and reactions are visualized as diamond-shaped nodes. The second part of the MPA_Pathway_Tool, the **“Pathway-Calculator”,** allows the **mapping of experimental data** on multiple pathways. Short videos for a detailed explanation about the MPA_Pathway_Tool are in progress.

### 1. Pathway-Creator

a) **Adding single reactions from KEGG- database**

If you want to add a reaction from the KEGG database you can click “KEGG REACTION”. Then you need to initialize your pathway by typing in your desired substrate and submitting. You will receive a list of potential products and can submit your desired product. After submitting your product you can choose the reaction. By clicking outside the window you can close it and the created nodes can be rearranged by drag and drop. For adding a new reaction from the KEGG- database, you can simply click on the compound- node, which you want to select as a substrate, with your left mouse button. After a window has emerged, you can submit the new product and new reaction.

b) **Adding a user-defined reaction**

If you can’t find a reaction in the KEGG database you can create your own reactions by clicking on “USER-DEFINED REACTIONS”. After another window emerges you can choose a substrate with a stoichiometric coefficient and add it to the reaction by clicking “ADD SUBSTRATE”. Additional substrates can be added by choosing the next substrate and stoichiometric coefficients and clicking again “ADD SUBSTRATE”. The same procedure can be performed with products. Then you have to set a reaction name. Additional information like K- numbers, EC- numbers and taxonomic requirements can be added by typing the information in the respective text field one by another. Finally you can see all submitted reaction information by clicking “SHOW REACTION” and submit the reaction by clicking “SUBMIT”. The window can be closed by clicking outside the window. The created nodes can be rearranged by drag and drop.

c) **Importing multiple reactions**

You have various possibilities for importing multiple reactions at once. At first you can upload a complete pathway in the “UPLOAD”- section as an SBML-file. The second possibility is to import a KEGG- Module by clicking on “IMPORT MULTIPLE REACTIONS”, “KEGG-MODULE”, searching your desired pathway and submitting. You can also import multiple reactions by EC- numbers by clicking “IMPORT MULTIPLE REACTIONS”, “IMPORT BY EC-NUMBERS” and either setting multiple EC- numbers (format: “1.1.1.1;1.1.1.2”) or adding iteratively adding single EC-numbers by choosing the desired EC-numbers and clicking “ADD EC NUMBER”. By clicking submit you will receive all reactions from the KEGG-database connected to the respective EC-number. You can select then your reaction and submit this reaction. The same procedure works for K-numbers as well. The format for setting multiple K-numbers are “K00001;K00002”.

d) **Modification of nodes**

You can delete an existent node by clicking on it with the right mouse button. By double clicking with the left mouse button on a node, a new window containing the identifier of the nodes emerges. It also allows changing the opacity under “key- Compound?” of the node by either selecting “YES” (setting the opacity to 100 %) or “NO” (setting the opacity to 40%). If you have clicked on a reaction node, you have access to further features. You can reverse the direction of the arrows by clicking on “REVERT DIRECTION”. You can also switch between reversibility and irreversibility of the reaction by clicking on the Checkbox for “Reversible”. If reversible is selected, arrow heads will point in both directions. If it is not selected, arrow heads will only point in one direction. Finally you can add a taxonomic requirement by selecting the taxonomic rank, typing in your desired taxonomy and clicking “ADD TAXONOMY”. Selected taxonomic requirements can be deleted by clicking on the respective delete- icon.

e) **Experimental data**

After uploading experimental data under “UPLOAD” and “Upload experimental data”, on the bottom emerges buttons for selecting an individual sample of the experimental data and a heatmap for adjusting the colors. Users can choose a sample, and as a consequence, the abundance of each reaction is shown by a specific color. By clicking on a reaction- node with the left mouse button a new window emerges containing information like the reaction name, K- numbers, EC- numbers and a heatmap with all features (e.g., proteins) mapped on the selected reaction for each sample. The heatmap on the bottom allows adjusting the colors. The mapped experimental data can be downloaded as CSV under “DOWNLOAD” and “DOWNLOAD HEATMAP”.

f) **Modification of the pathway**

Under the button “NODE CONFIGURATIONS” you have multiple options for modify your pathway. At first you enable “force”. This leads to attraction and repulsion of nodes based on their connections. You can also set abbreviations by clicking “ABBREVIATIONS”. After a new window has opened you can type in an abbreviation in the respective text field and then submit this abbreviation. You can close the window but clicking outside the window. You can also split one node into separate nodes. After clicking “SPLIT NODES” a new window emerges, where you can add compounds. After submitting, each node is split into separate nodes with only one link. Merging nodes into a single node is possible by clicking on “MERGE NODES”, adding the nodes you want to merge, choosing and submitting a name for the merged nodes and at the end clicking “MERGE SELECTED NODES”. You can add taxonomic requirements for each reaction under “ADD TAXONOMY” for the complete pathway or for individual reactions. You can close the window but clicking outside the window.

g) **Download and upload**

Created pathways can be downloaded as CSV, SVG, JSON, and SBML by clicking on the button “DOWNLOAD” and the respective exchange format. You can download the stoichiometric matrix of your metabolic pathway as CSV by clicking “DOWNLOAD” and “STOICHIOMETRIC MATRIX”.

Already created pathways can be reused by clicking “UPLOAD” and then upload the respective pathway as CSV (“Upload pathway as CSV”), JSON (“Upload pathway as JSON”), and SBML (“Upload pathway as SBML”).

### 2. Pathway-Calculator

If you want to map experimental data on multiple pathways automatically you can use the “Pathway-Calculator”. On the left, you can upload your experimental data (see https://github.com/danielwalke/MPA_Pathway_Tool/tree/main/experimental%20data%20example for an example file). On the right, you can add your created pathways as CSV, JSON or SBML. After selecting all files, you can click “START”. After experimental data have been mapped on each pathway, you can download the mapped data as CSV. You can also download the unmatched features (e.g., unmatched proteins).
